# SCIΦ: Single-cell mutation identification via phylogenetic inference

**DOI:** 10.1101/290908

**Authors:** Jochen Singer, Jack Kuipers, Katharina Jahn, Niko Beerenwinkel

**Affiliations:** Department of Biosystems Science and Engineering, ETH Zurich, Basel, Switzerland; SIB Swiss Institute of Bioinformatics, Basel, Switzerland

**Author notes:** These authors contributed equally (Jochen Singer, Jack Kuipers).

## Abstract

Understanding the evolution of cancer is important for the development of appropriate cancer therapies. The task is challenging because tumors evolve as heterogeneous cell populations with an unknown number of genetically distinct subclones of varying frequencies. Conventional approaches based on bulk sequencing are limited in addressing this challenge as clones cannot be observed directly. Single-cell sequencing holds the promise of resolving the heterogeneity of tumors; however, it has its own challenges including elevated error rates, allelic dropout, and uneven coverage. Here, we develop a new approach to mutation detection in individual tumor cells by leveraging the evolutionary relationship among cells. Our method, called SCIΦ, jointly calls mutations in individual cells and estimates the tumor phylogeny among these cells. Employing a Markov Chain Monte Carlo scheme we robustly account for the various sources of noise in single-cell sequencing data. Our approach enables us to reliably call mutations in each single cell even in experiments with high dropout rates and missing data. We show that SCIΦ outperforms existing methods on simulated data and applied it to different real-world datasets, namely a whole exome breast cancer as well as a panel acute lymphoblastic leukemia dataset. Availability: https://github.com/cbg-ethz/SCIPhI

## 1 Introduction

Due to recent technological advances it is now possible to sequence the genome of individual cells [20]. This allows, for the first time, to directly study genetic cell-to-cell variability and gives unprecedented insights into somatic cell evolution in development and disease.

Having single-cell resolution is especially useful for the analysis of intra-tumor heterogeneity [19]. This is due to the central role that mutational heterogeneity and subclonal tumor composition play in the failure of targeted cancer therapies, where resistant subclones can initiate tumor recurrence [2, 8]. Presently, genetic analyses of tumors are mostly based on sequencing bulk samples which only provides admixed variant allele frequency profiles of many thousands to millions of cells. These aggregate measurements are, however, only of limited use for the inference of subclonal genotypes and and their phylogenetic relationships [10, 12]. The two main issues are that mutational signals of small subclones can not be distinguished from noise and that the deconvolution of the aggregate measurements into clones is, in general, an underdetermined problem.

In contrast, single-cell sequencing data provides direct measurements of cellular genotypes thus bypassing the deconvolution problem of bulk measurements. However, this advantage comes at the cost of elevated noise due to the limited amount of DNA material present in a cell and the extensive DNA amplification required prior to sequencing. The most common approach for this initial amplification of single-cell DNA is Multiple Displacement Amplification (MDA) [14]. While this process is very efficient at amplifying the overall DNA material, high rates of allelic drop-out, i.e., the random non-amplification of one allele of a heterozygous genotupe site, are observed. Starting with the DNA of a single cell, all evidence of a heterozygous genotype mutation is lost when the mutated allele drops out, which happens at a rate of about 10% to 20%. Also, false positive artifacts can arise in the MDA amplification when random errors introduced early in the process end up with high frequencies due to allelic amplification biases. Further challenges arise from uneven amplification across the genome which results in non-uniform coverage that will leave some sites with insufficient coverage depth for reliable base calling.

These technical issues result in single-cell-specific noise profiles for which regular variant callers developed for next-generation sequencing data, such as the Genome Analysis Toolkit (GATK) HaplotypeCaller [18] or SAMtools [17], are ill-suited. Two single-cell specific mutation callers, namely Monovar [25] and SCcaller [4], have therefore been recently developed. Both methods take raw sequencing data (BAM files) and output the inferred genotypes of the cells. Monovar specifically addresses the problem of low and uneven coverage in mutation calling by pooling sequencing information across cells, while assuming that no dependencies exist across sites. In contrast, SCcaller detects variants independently for each cell and accounts for local allelic amplification biases. However, the identification of such biases is based on germline single-nucleotide polymorphisms (SNPs), which might not be available, for example, for panel sequencing data. Further, it cannot recover mutations from dropout events or loss of heterozygosity.

Here, we present SCIΦ, a new single-cell-specific variant caller that combines single-cell genotyping with reconstruction of the cell lineage tree. SCIΦ leverages the fact that the somatic cells of an organism are related via a phylogenetic tree where mutations are propagated along tree branches. SCIΦ can reliably identify single-nucleotide variants (SNVs) in single cells with very low or even no variant allele support. We show that SCIΦ outperforms Monovar, the only other tool able to transfers information between cells, on simulated and real data.

## 2 Methods

Our inference scheme starts with an initial identification of possible mutation loci and then performs joint phylogenetic inference and variant calling via posterior sampling. After introducing the general model for nucleotide frequencies, we describe these steps in more detail.

### 2.1 Nucleotide frequency model

We model the nucleotide counts *s* at a locus with total coverage *c* using the beta-binomial distribution [e.g., 7, 23] as

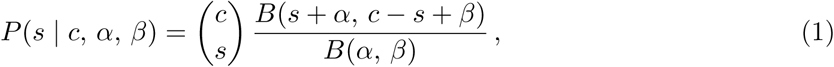

with parameters *α* and *β* and where *B* is the beta function. For our implementation we will employ an alternative parametrization of the beta binomial distribution with 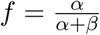 being the frequency of a nucleotide and *ω* = *α* + *β* an overdispersion term which decreases with increasing variance.

For locus *i* and cell *j* with coverage *c*_*ij*_, the probability of the observed count (support) *s*_*ij*_ for a specific nucleotide in the absence of a mutation is

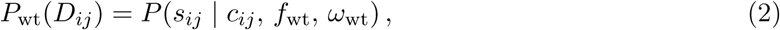

where *D*_*ij*_ = (*s*_*ij*_, *c*_*ij*_) and *f*_wt_ is the expected frequency of the observed nucleotide, which, for example, could have arisen from sequencing error. Large values of *ω*_wt_ lead to a binomial distribution representing independent sequencing errors. In the presence of a heterozygous mutation, the probability of the counts is

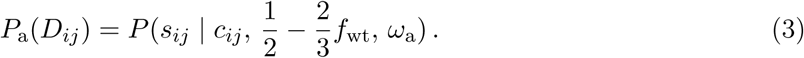

The underlying allele frequency of 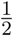 is corrected by sequencing errors producing any of the other two bases. Low values of the overdispersion term *ω*_a_ reflect a small number of initial genomic fragments and any additional feedback in the amplification. SCIΦ assumes copy number neutrality, but learning *ω*_a_ allows for compensating for shifts in the mean variant allele frequency away from 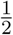.

### 2.2 Identification of candidate mutated loci

Likely mutated loci are identified using the posterior probability of observing at least one mutated cell at a specific locus. The probability of observing no mutation at locus *i* across all cells is

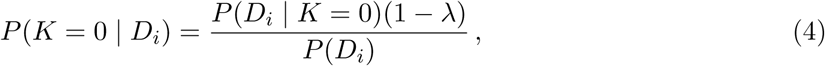

where *K* is a random variable indicating the number of mutated cells and *λ* is the prior probability of a mutation occuring at the locus. The probability of observing the mutation in *k* cells is

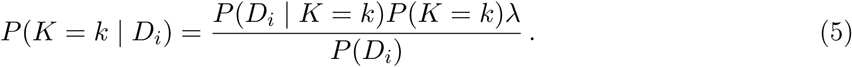

We do not need to compute *P* (*D*_*i*_) as it cancels out when computing the likelihood ratio or posterior odds.

The likelihood of the data given that exactly *k* of the *m* cells possess the mutation, is given by

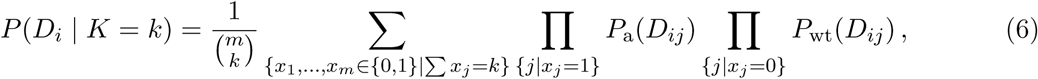

where *x*_*i*_ indicates whether cell *i* is mutated or not. *P* (*D*_*i*_ | *K* = *k*) can be computed efficiently using a dynamic programming approach [as in 25, 15].

The prior probability of a mutation in a phylogeny affecting *k* descendant cells is determined by placing mutations uniformly among the edges of the tree (Section A.1) leading to

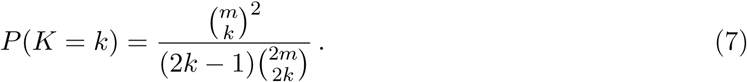

### 2.3 Allelic dropout

Along with the uncertainty in the supporting read counts due to the amplifications in each cell when a mutation is present, an additional artifact is dropout whereby one allele is not amplified at all. To account for allelic dropout occuring with probability *µ*, we introduce the following mixture for the likelihood of the observations for each cell:

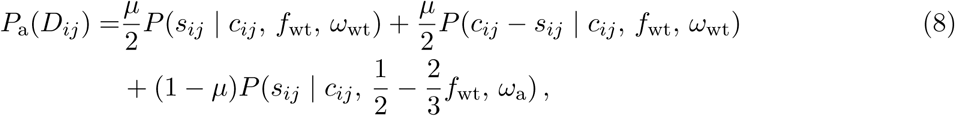

where the first term describes the loss of the mutant allele, the second the loss of the wild-type allele and the third term describes a heterozygous mutation. The case *µ* = 0 reduces to Equation (3).

### 2.4 Tree likelihood

Our model to infer tumor phylogeny consists of three parts [akin to 11]: the tree structure *T*, the mutation attachments to edges *σ*, and the parameters of the model *θ* (the parameters *f*_*wt*_, *ω*_wt_, and *ω*_a_ previously introduced, the dropout mixture coefficient *µ* as well as a homozygosity coefficient which we will introduce later). We represent the phylogeny of a tumor using a genealogical tree. Here the *m* sampled tumor cells are represented by leaves in a binary tree and the mutations are placed along the edges. There are (2*m* − 3)!! different tree structures, while each of the *n* mutations can be attached to the (2*m* − 1) edges leading to (2*m* − 3)!!(2*m* − 1)^*n*^ possible configurations for the discrete component (*T, σ*) of our model. As a result, it is infeasible to enumerate all solutions. Instead we employ a Markov Chain Monte Carlo approach to search and sample from the tree space.

In order to do so, we employ the likelihood of a specific tree realization with the mutation attachment parameter *σ* and the parameters *θ* to be

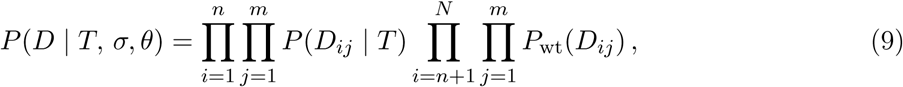

where *P* (*D*_*ij*_ | *T*) = *P*_a_(*D*_*ij*_) if the cell *j* is below mutation *i* (on the path from leaf *j* to the root) and *P* (*D*_*ij*_ | *T*) = *P*_wt_(*D*_*ij*_) otherwise. The first set of products describes the loci identified to be likely mutated (Section 2.2) which are placed on the tree and used to infer its phylogenetic structure. The second half represents all loci where no mutation is present which inform the inference of the sequencing error parameters.

Analogously to [11] we marginalize out the attachment points of the mutations. Assuming each mutation is equally likely to attach to any edge in the tree and the attachment probability to be independent between mutations we have 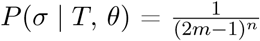 so that

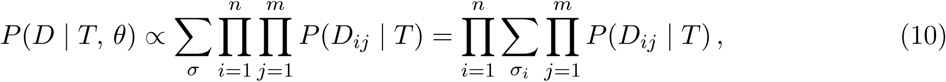

For each locus, the sum can be written explicitly as

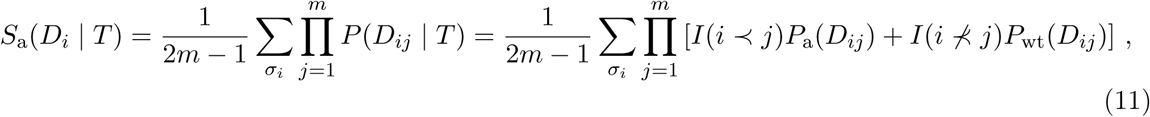

where *I* is the indicator function and (*i* ≺ *j*) indicates that cell *j* sits below the attachment point *σ*_*i*_ of mutation *i* in the tree *T*. The sum can be computed in *O*(*m*) time using the binary tree structure. Employing *T* we propagate the probability of attaching a mutation to a specific node from the leaves towards the root. This can be implemented using the depth first search (DFS) algorithm, combining in each node the probabilities from two previously computed subtrees.

Computing Equation (10) is therefore in *O*(*mn*) while the marginalisation has the further benefit of reducing the search space by a factor of (2*m* − 1)^*n*^. In addition we employ the marginalisation to focus on the tree structure of the cell lineage rather than the annotation with mutations.

Making use of the factorization of the beta-binomial density function into Gamma functions, the term 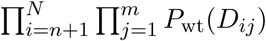 in Equation (9) can be computed in time linear in the number of different coverages of the sequencing experiment (Section A.2). Since that number is typically much smaller than *mn* for sequencing projects, the overall runtime is dominated by *O*(*mn*).

### 2.5 Accounting for zygosity

Because tumor cells show chromosomal abnormalities, mutations can be observed as homozygous variants even without dropout events. In order to also account for loss of heterozygosity, we adapt the scheme introduced in Section 2.4. Instead of computing the likelihood of the data when attaching a mutation to a node in the lineage tree in the heterozygous state only, we additionally compute the likelihood when attaching each mutation in the homozygous state, and define the sum

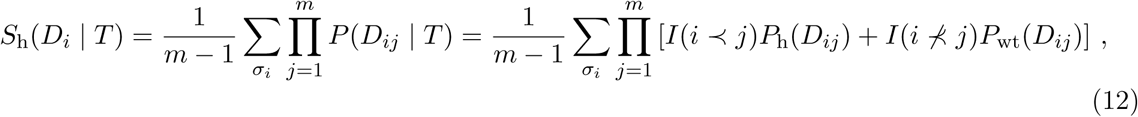

involving the nucleotide model when only alternative alleles are present

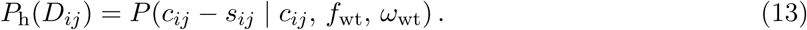

Note that homozygous mutations are only attached to inner nodes as the probability of observing a dropout event in a single cell is assumed to be higher than a single homozygous mutation.

Utilising the tree structure, the sum can again be computed in *O*(*m*) time for each mutation on the tree. The overall likelihood for each mutation is a weighted sum of the two possibilities leading to

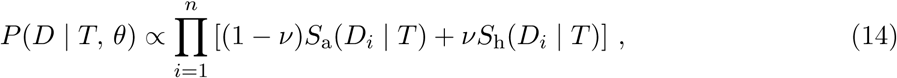

with homozygosity coefficient *ν*. Thus, we allow certain violations of the infinite sites assumption [13] by capturing homozygous mutations which are not due to dropout events.

### 2.6 Markov Chain Monte Carlo sampling

Using the tree likelihood we employ an MCMC scheme to sample from the posterior distribution of mutation assignments as well as tree structures given the data (for simplicity with uniform priors). In order to do so, we propose a new state (*T* ′, *θ*′) from the current state (*T, θ*) making use of properly defined moves such that the chain is ergodic. We change one parameter at a time with transition probability *q*(*T* ′, *θ*′ | *T, θ*) and accept the new configuration with probability

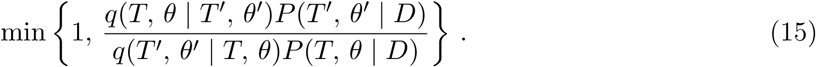

The tree structure can be changed using the *prune and reattach* move. Here we randomly draw a node from the tree and re-attach it to a random node not contained in the pruned subtree. This move is reversible, irreducible, and aperiodic. Additionally we include a move which swaps two leaf nodes. For the parameters of the beta-binomial distribution, the dropout coefficient *µ* (and the homozygosity coefficient *ν*) we perform independent random Gaussian walks. The standard deviations of the steps are adjusted using adaptive MCMC [1] to track an acceptance rate of 50%.

We sample proportional to *P* (*T, θ* | *D*) from the posterior distribution after a burn-in phase. Convergence is achieved after *x* iterations, with heuristic arguments suggesting *x* ∝ *m*^2^ log(*m*) [11]. The overall runtime complexity is *O*(*x* × max(*mn, c*)) with *c* being the number of unique coverage values of the experiment. From the sample of trees and parameters we could also conditionally sample the placement of the mutations for the full joint posterior sample. Instead, utilising the full weights of attaching each mutation to different edges we record the probability of each cell possessing each mutation. Averaging over the MCMC chain provides the posterior genotype matrix and hence our single-cell variant calls.

### 2.7 Simulation of ground truth datasets

In order to benchmark the performance of SCIΦ we simulated tumor evolution by introducing a cell lineage tree and simulated read counts by mimicking the noisy MDA process. For *m* cells, we created a random binary genealogical cell linage tree with 100 mutations attached to the edges. The placement of the mutations defines which cells possess each mutation and was chosen such that each mutation is shared by least two cells. Further among all the mutations present in cells a specified fraction *µ* was randomly selected as dropouts, i.e. 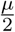 of the mutations became wild type and 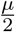 became homozygous.

Then we generated an artificial reference chromosome of 1 million base pairs (bp) and divided it into segments of approximately 1000bp for each cell individually. For these segments, we generated a coverage distribution following a negative binomial distribution with a mean of 25 nucleotides and a variance of 50. Additionally, 10% of the segments were assigned 0 coverage to include missing information. The coverage of specific positions was additionally randomized following a discretized Gaussian distribution with the segment coverage as mean and a standard deviation of 10% of that mean in order to simulate the uneven coverage profiles of real single-cell sequencing experiments.

For simulating nucleotides under the MDA process, we drew them from a Pólya urn model. While heterozygous positions contain two chromosomes, one with two wild type strands and one with two mutant strands (*α* = 2, *β* = 2), dropout positions only retain the two strands of one of the two chromosomes (*α* = 2, *β* = 0; *α* = 0, *β* = 2,). A strand is then randomly chosen, copied, and returned to the urn together with the copy. With a probability of 5 × 10^*-*7^ the copy will be mutated and an allele different from the original one is returned, corresponding to the error rate of the MDA polymerase (10^*-*6^ − 10^*-*7^ [3]). This process is repeated *c* times and the copies are retained. Finally, with probability of 5 × 10^*-*4^, a nucleotide is mutated to account for sequencing errors, and the resulting simulated data was embedded into a multi-pileup file. Additional simulations are reported in Section A.4.

## 3 Results and Discussion

In order to investigate the performance of SCIΦ we conducted several experiments on simulated data and additionally on several real datasets. We compared SCIΦ to Monovar [25], the only published single-cell mutation caller sharing information across cells. We start by analyzing the results of the simulated data.

### 3.1 Benchmarks for simulated data

We first investigated how the performance depends on the number of cells sequenced in the experiment. SCIΦ is more sensitive in calling mutations than Monovar while showing comparable precision in all settings analyzed (Figure 1). The reason for this is twofold: First, due to the tree inference SCIΦ can assign a mutation to a particular cell with very low or even missing variant support at a specific locus. Second, making use of a beta-binomial model to represent the nucleotide counts and learning its parameters accurately reflects the underlying process generating nucleotide counts.

**Figure 1:**
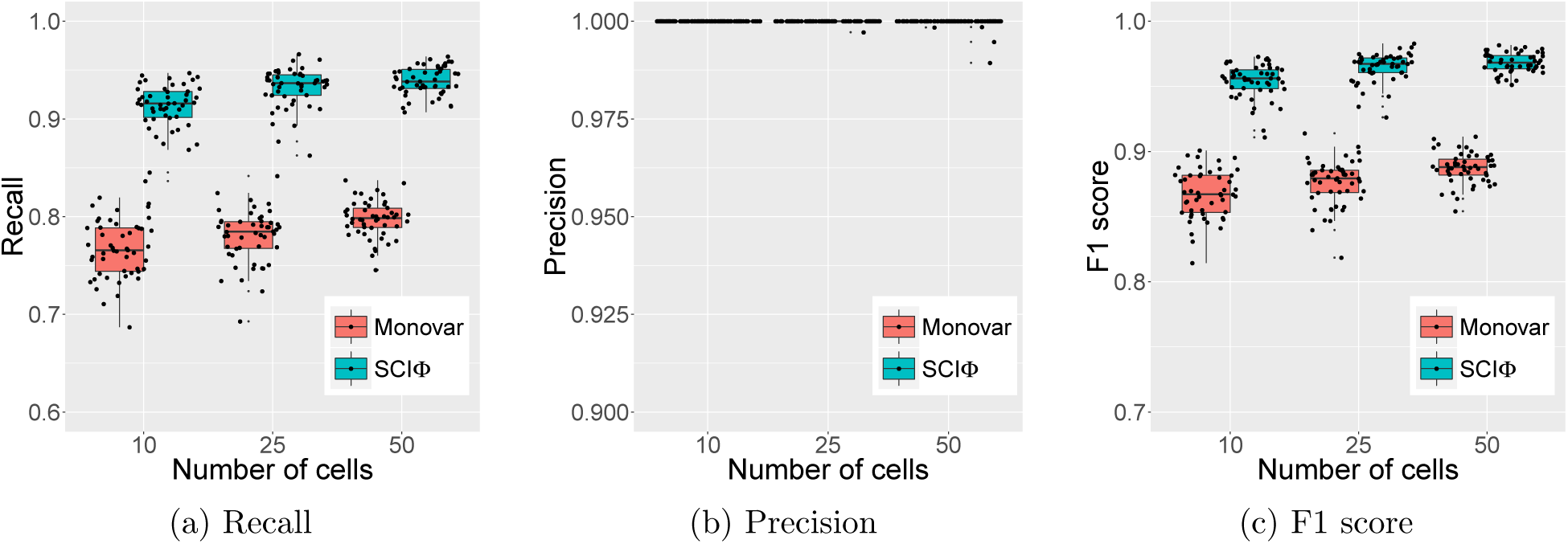
Summary statistics of the different performance measures from SCIΦ and Monovar on simulated data with different number of cells.

Due to the observed large range of dropout rates, ranging from 10% to more than 40% [12], a second experiment was conducted to explore the dependence of the methods on the dropout rate of the experiment. Here we concentrated on dropout rates of 10, 20, and 30%. Since the exact dropout rate of a dataset is often not known, we used the default values of the callers, namely 20% for Monovar and 10% for SCIΦ (Figure 2a). Note that SCIΦ learns the dropout rate and uses 10% only as a starting condition.

**Figure 2:**
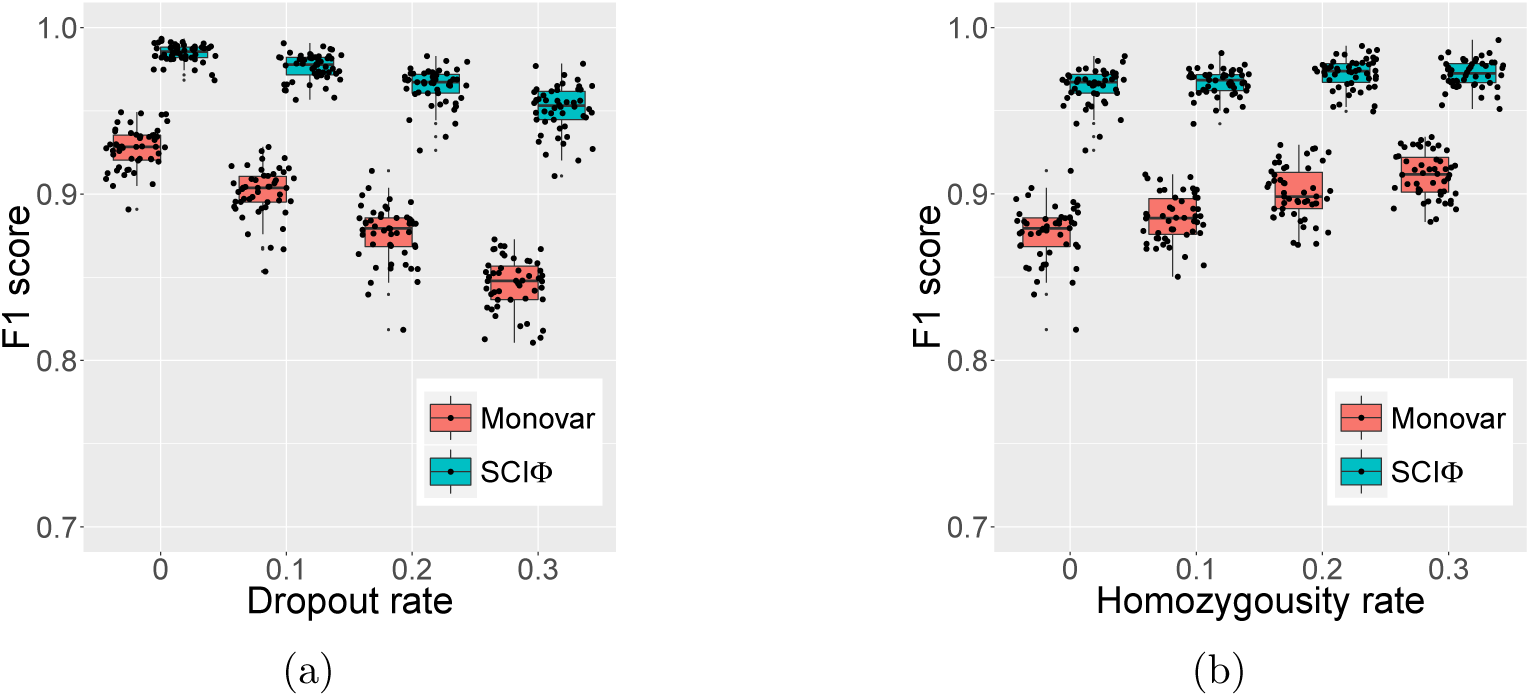
Summary statistics of the F1 performance measure from SCIΦ and Monovar on simulated data with different levels of dropout events (a) and homozygosity rates (b).

We found SCIΦ to be more robust to increasing dropout rates in comparison to Monovar (Figure 2a). In addition to using the phylogentic tree structure, SCIΦ also learns the dropout rate of the experiment during the MCMC scheme.

An additional experiment was conducted to investigate the effects of loss of heterozygosity. Monovar as well as SCIΦ perform better with increasing levels of homozygous mutations present in the experiment (Figure 2b). Especially Monovar benefits from homozygous mutations as these are very unlikely to be classified as wild type. SCIΦ experiences a more modest benefit from homozygous mutations since it already starts with high performance due to the usage of the phylogenetic tree structure to accurately call mutations.

### 3.2 Application to real data

We applied SCIΦ to two human tumor sequencing datasets. The first dataset is described in [24] (Sequence Read Archive (SRA) accession numbers SRA05319), where the authors performed exome sequencing on single and bulk cells of a breast cancer patient. Here we identified somatic mutations in 16 single cells using bulk sequenced normal control dataset to distinguish somatic from germline mutations (see Section A.3 for details). This dataset is particularly challenging because cells are aneuploid.

We identified around 50% of the mutations to be shared across all cells and therefore placed them into the root of the inferred phylogenetic tree (Figure 3a). The assignment of different mutations to different subclones is depicted in Figure 3a. For example, 263 mutations distinguish cell *h1* from the other cells and 200 mutations separate the lineage of cells *a1, a4*, and *a6* from the remaining tree. The posterior probabilities of each cell possessing each mutation show the grouping into subclones (Figure 3b). Using the tree inferred by SCIΦ to order the mutation calls of Monovar (Figure 3c) allows a more direct comparison. The assignment of mutations to cells is very homogeneous for the subclones using SCIΦ (Figure 3b). In contrast, the mutation assignment based on Monovar’s inferred probabilities is much more noisy (Figure 3c).

**Figure 3:**
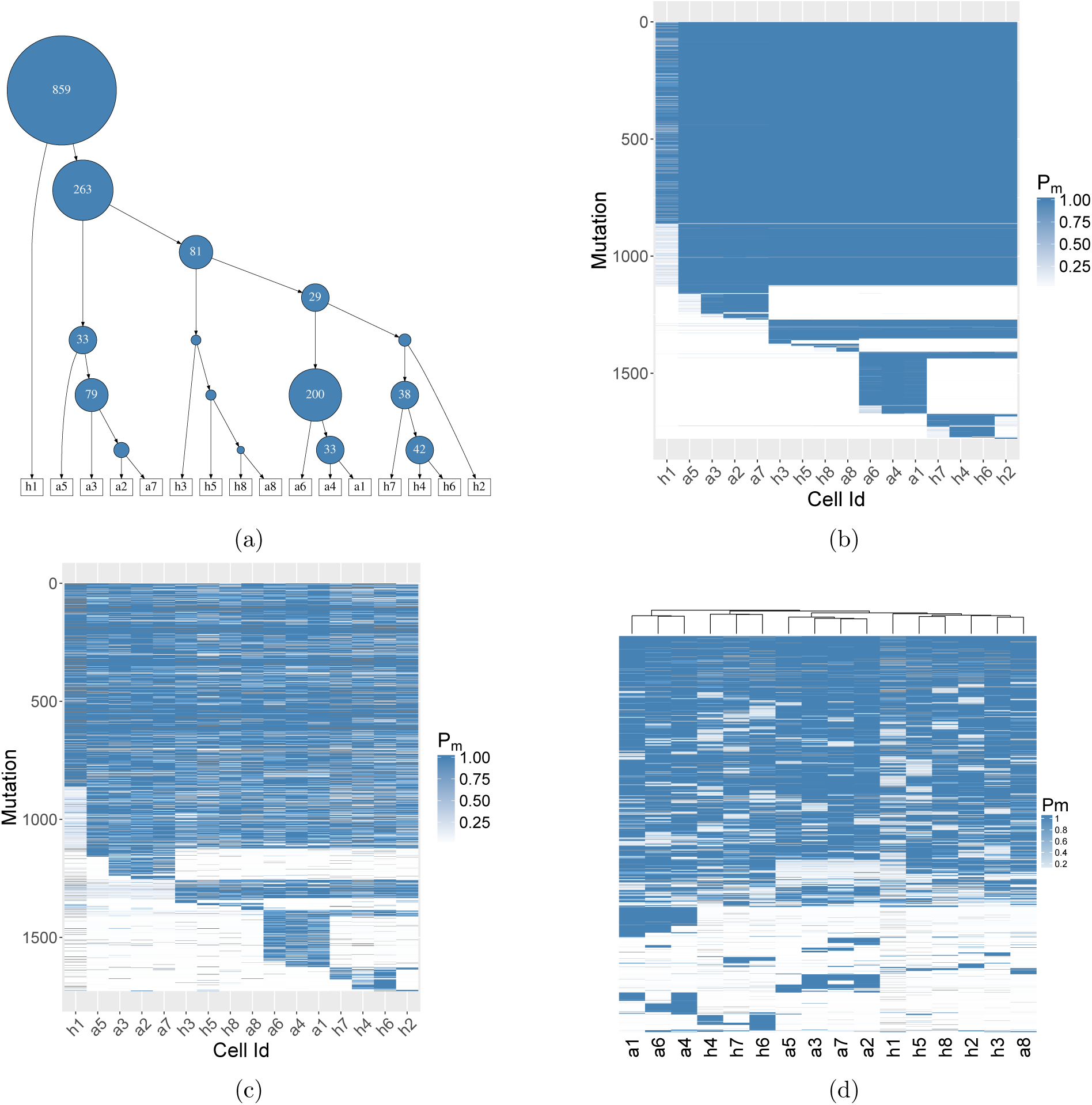
Summary of the mutation calls obtained with Monovar and SCIΦ on a breast cancer patient dataset [24] consisting of 16 single tumor cells and a control normal bulk sequencing dataset. (a) Cell lineage tree with mutation attachment identified by SCIΦ. The area of a node is proportional to its number of assigned mutations. (b) Posterior probability of SCIΦ mutation calls clustered according to the tree in a). (c) Probability of Monovar mutation calls for loci identified as mutated by SCIΦ and clustered according to the tree in a). (d) Probability of Monovar mutation calls for loci identified as mutated by SCIΦ and clustered hierarchically.

It is interesting to observe that Monovar identifies additional mutations for cell *h*1 above mutation index 1000 (Figure 3c). Investigating these mutations more closely shows that they all have very low coverage (1–4 reads) and in most cases no alternative nucleotide support. Further, some of these mutations with coverage 1 and no alternative support were labeled homozygous alternative, which is unlikely as this would require a back mutation or sequencing errors to the reference allele for several mutations.

Without using the tree inferred by SCIΦ to order the mutation calls from Monovar, hierarchal clustering (Figure 3d) leads to a similar subclonal structure to SCIΦ (Figure 3b). However, there are some differences. For example, *h2* is hierarchally clustered with *h5, h8, h3* and *a8*, rather than with *h4, h6, h7* and *h8*. The hierarchal clustering does not enforce a phylogenetic tree and weights false negative and false positive signals equally. However, from SCIΦ (Figure 3a) we can see that cell *h2* is only missing mutations which are in common in cells *h4, h6, h7*. Therefore, its placement earlier in the tree above those cells is much more evolutionarily plausible.

The second dataset consists of 255 cells from a patient (number 3) with acute lymphoblastic leukemia sequenced using a panel sequencing approach [6] (SRA accession number SRP044380). The results (Figure 4) highlight similar aspects to those mentioned for the previous breast cancer dataset, especially the much less noisy mutation assignment. It is interesting to observe that SCIΦ not only recovered dropouts, but also assigned much lower mutation probabilities to likely wild type positions compared to Monovar (Figure 4).

**Figure 4:**
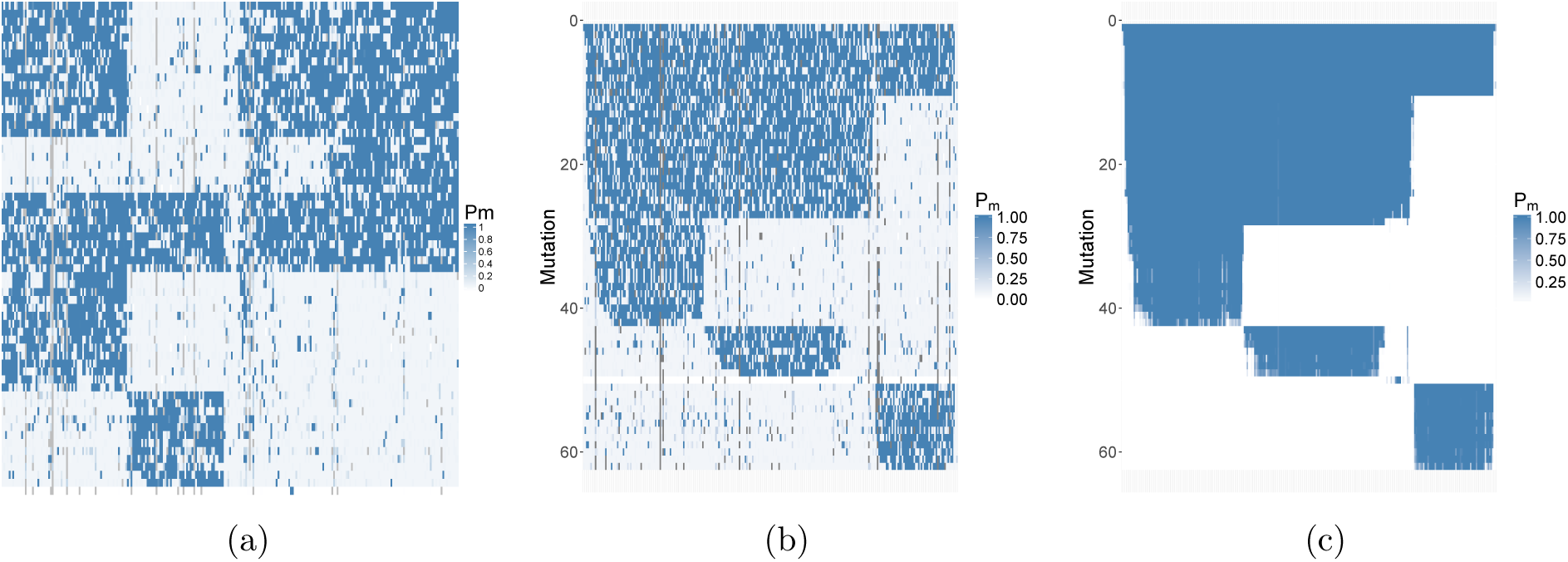
Summary of the mutation calls from SCIΦ and Monovar on a dataset consisting of 255 cells from a patient (number 3) with acute lymphoblastic leukemia [6]. (a) Monovar mutation calls for loci identified as mutated by SCIΦ clustered hierarchically. (b) Monovar mutation calls clustered according to the tree inferred by SCIΦ. (c) SCIΦ mutation calls clustered according to its inferred tree.

## 4 Conclusions

Single-cell sequencing allows us to directly study genetic cell-to-cell variability and gives unprecedented insights into somatic cell evolution. This is of particular interest in cancer genomics because tumors show heterogeneous cell compositions often resulting in the failure of targeted cancer therapies. Here, we introduced SCIΦ, the first single-cell mutation caller that simultaneously infers the mutational landscape and the phylogenetic history of a tumor sample. SCIΦ accounts for the elevated noise levels of single cell data by appropriately modeling the genomic amplification process and the high fraction of dropout events. In combination with a Markov Chain Monte Carlo phylogenetic tree inference scheme, mutations are reliably assigned to individual cells.

We have compared SCIΦ to Monovar [25] on both simulated and real datasets. For the simulated data, SCIΦ has comparable precision and significantly better recall and F1 score. For the real datasets, we showed that SCIΦ achieves a much cleaner assignment of mutations to cells within subclones. In particular, SCIΦ recovered mutations from dropout events using the inferred phylogenetic tree structure of the sample to share information across cells, whereas Monovar missed these events. Furthermore, the phylogenetic tree inferred by SCIΦ reflects the evolutionary history more accurately than a hierarchical clustering from Monovar. Mutation calling and lineage tree building are two interdependent tasks and addressing them in a single statistical model provides both improved mutation calls as well as a better estimate of the underlying cell lineage tree, and hence a better understanding of tumor heterogeneity.

## Acknowledgments

We would like to thank David Seifert for constructive discussions and C++ support as well as Franziska Singer for critical feedback.

## Software availability

SCIΦ has been implemented in C++ using [5] and is freely available under a GNU General Public License v3.0 license at https://github.com/cbg-ethz/SCIPhI.

## Appendix

### A.1 Probability of a mutation in *k* cells in a random binary tree

To compute the number of cells below a mutation in a binary genealogical tree, we work recursively. We record in *P*_*m*_(*k*) the probability of a mutation being placed uniformly among the edges of trees with *m* cells affecting exactly *k* of them. Moving from trees with *m* cells, which possess (2*m* − 1) edges, to trees with (*m* + 1) cells we can create a new internal node along any of the edges. This adds a further two edges. When doing so, the mutation may be above or below the new cell added, or placed along the two new edges leading to the recursion

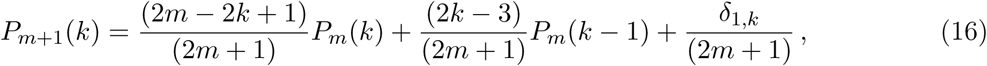

with initial condition of *P*_1_(1) = 1 and boundary conditions of *P*_*m*_(0) = 0. The solution to the recursion in Equation (16) is

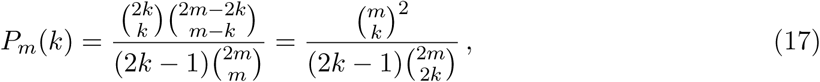

which can easily be shown by induction. We start with *k* = 1

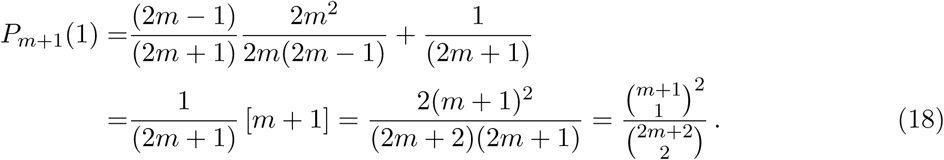

For 1 *< k ≤ m* we have

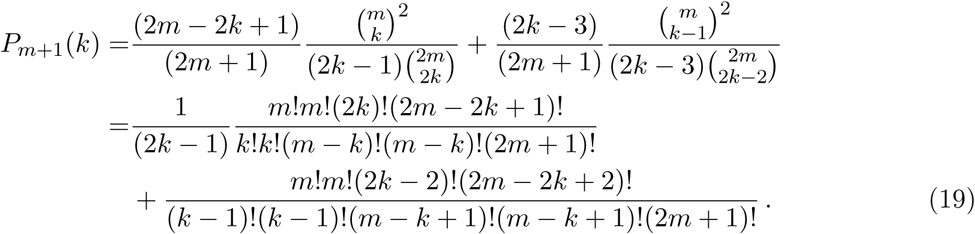

Taking out a common factor, this reduces to

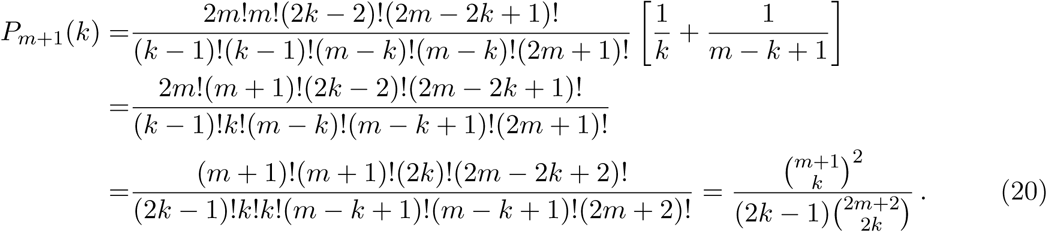

### A.2 Efficient likelihood computation for wild type loci

The beta-binomial density function of Equation (1) can be expressed using Gamma functions as:

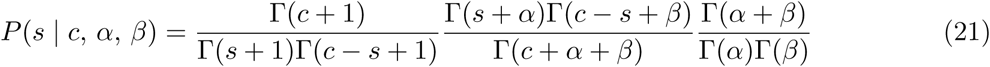

For numerical accuracy we compute log probabilities such that the factors become summands. These can be treated individually, leading to nine terms per locus. However, summands only involving *s* and *c* depend solely on the data and not on the tree or parameters. As they do not change between iterations of the algorithm, they can be ignored. The summands involving only *α* and *β* do not depend on the locus and can be computed in *O*(1) for all loci. This leaves us with three summands that need to be recomputed whenever one of the parameters of the model changes. For efficiency, we store an array with occurrences of unique values of *s, c* − *s* and *c*. This reduces the time complexity from *O*(*Nm*) to *O*(*c*) for the computation of the probability of all (*N* − *n*) wild type loci.

### A.3 Variant calling pipeline

In order to reliably call mutations, we applied several mapping and purification steps using NGS-pipe [22]. We first mapped the downloaded FASTQ files using BWA-mem [16] version 0.7.15 to the human reference genome *hg*19. The resulting files were merged, sorted and duplicates removed (for exome data) using Picard tools (http://broadinstitute.github.io/picard/) version 2.8.3. Afterwards we realigned the reads around indels using the GATK [18] version 3.5. SAMtools mpileup [17] version 1.3.1 was used with parameters `-A -B -d 1000 -q 40` for the exome dataset and `-A -B -d 100000 -q 40` for the panel dataset. Monovar (commit 7b47571) was then run according to the authors’ recommendation (https://bitbucket.org/hamimzafar/monovar). In order to compare the results, the normalized and Phred-scaled likelihoods for the genotypes reported by Monovar were back-transformed and the fraction of heterozygous plus homozygous genotype likelihoods reported as the probability of the mutation. Further, hierarchal clustering of Monovar mutation calls was performed in R [21] version 3.4.0 using *ComplexHeatmaps* [9]. SCIΦ was run with default values with an additional filter of requiring at least two cells to show an alternative nucleotide count of at least three and a prior mutation rate of 0.001.

Further, variants with low VAFs across all cells for a given locus where filtered out by SCIΦ using a likelihood ratio test. In order to do so, we first fitted a beta-binomial distribution with free mean and overdispersion to maximise the likelihood of that locus across all cells with non-zero variant reads and computed the likelihood *L*_1_. For the same set of cells, a second beta-binomial distribution with a mean fixed to 0.25 (to allow for copy number changes) and free overdispersion was fitted to compute the maximum likelihood *L*_0_ of the constrained model. The test statistic 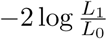 is asymptotically 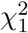 distributed and loci with a p-value > 0.05 or estimated mean > 0.25 were kept. For the exome data, loci with coverage less than six in the control bulk sequencing dataset were excluded as not reliably distinguishable from germline variants. Further, loci showing an alternative nucleotide count of 2 or more in the bulk control were excluded as germline variants.

### A.4 Influence of prior parameters

In an additional experiment we investigated the influence of changing the prior parameters for SCIΦ and Monovar (Figure 6). For SCIΦ we changed the prior probability *λ* of a locus to be mutated, where the default is 0.0001. For Monovar we investigated the influence of the prior probability of a false-positive error.

**Figure 5:**
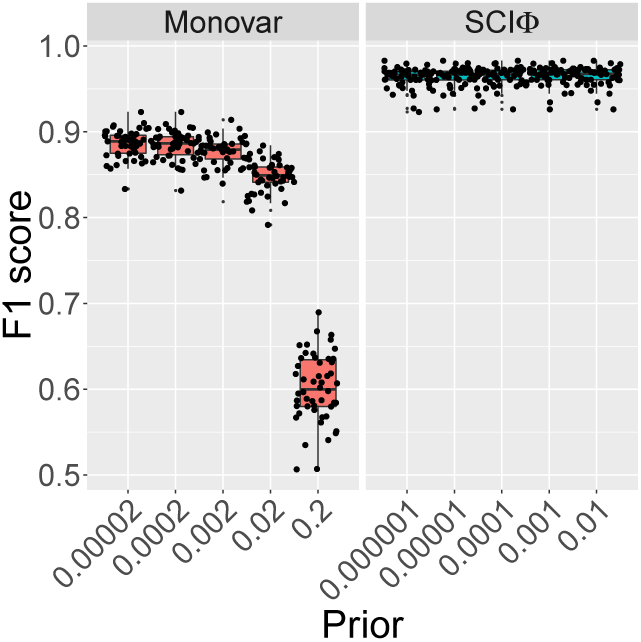
F1 score

The F1 score does not change dramatically over five orders of magnitude (except with a very high prior for Monovar).

### A.5 Results for alternative simulation

As can be observed in Figure 7 the nucleotide frequency distribution after the MDA does not necessary follow a beta binomial distribution with *α* and *β* of the beta distribution equal to 2. Instead, here the distribution is uniform with the exception of frequencies of 0 and 1, which correspond to no mutation observed or the mutation being present in homozygous state. This scenario can be simulated with a beta binomial distribution where *α* and *β* are set to 1. Therefore, we provide additional benchmarks on simulated data following the same scheme as in 2.7 with *α* and *β* set to 1 for a heterozygous genotype and *α* set to 1 and *β* set to 0 (or vice versa) for the homozygous reference or homozygous alternative genotype. The results are similar to the previous simulation and summarized in Figure 8.

Figure 6: Summary statistics of F1 performance from SCIΦ and Monovar on simulated data as their prior parameters are varied.

**Figure 7:**
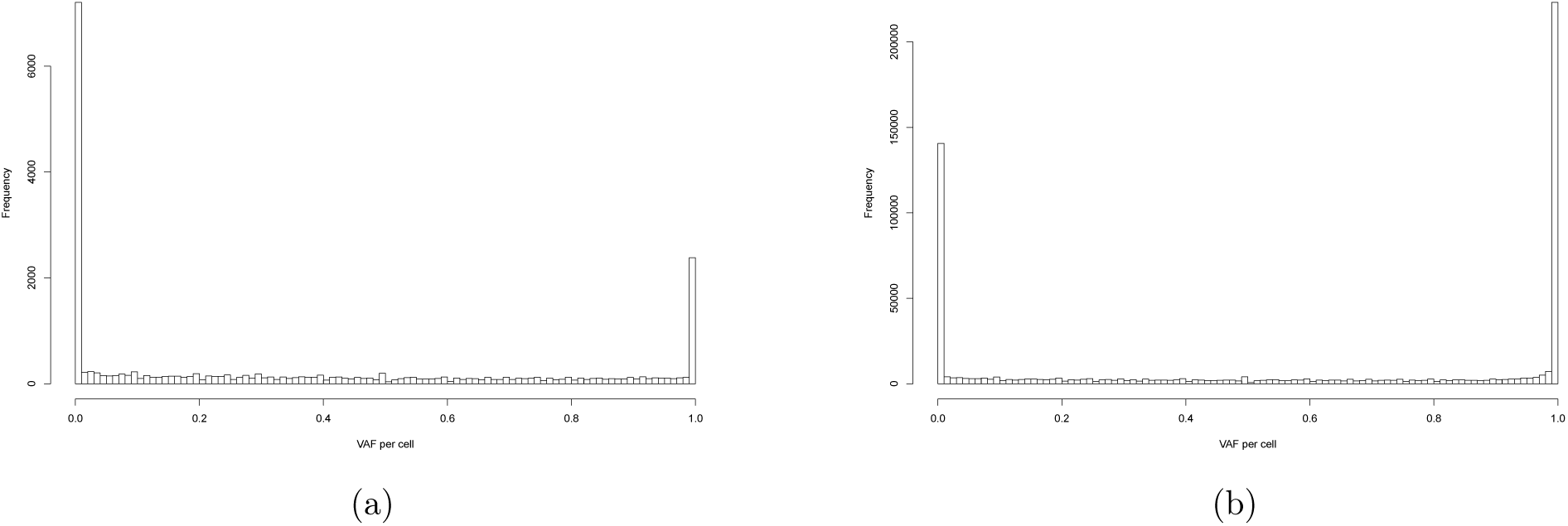
Variant allele frequencies (VAFs) of the single cells for positions containing somatic mutations identified by SCIΦ (a) and positions with differences to the reference genome identified by Monovar (b) in the data described in [24].

**Figure 8:**
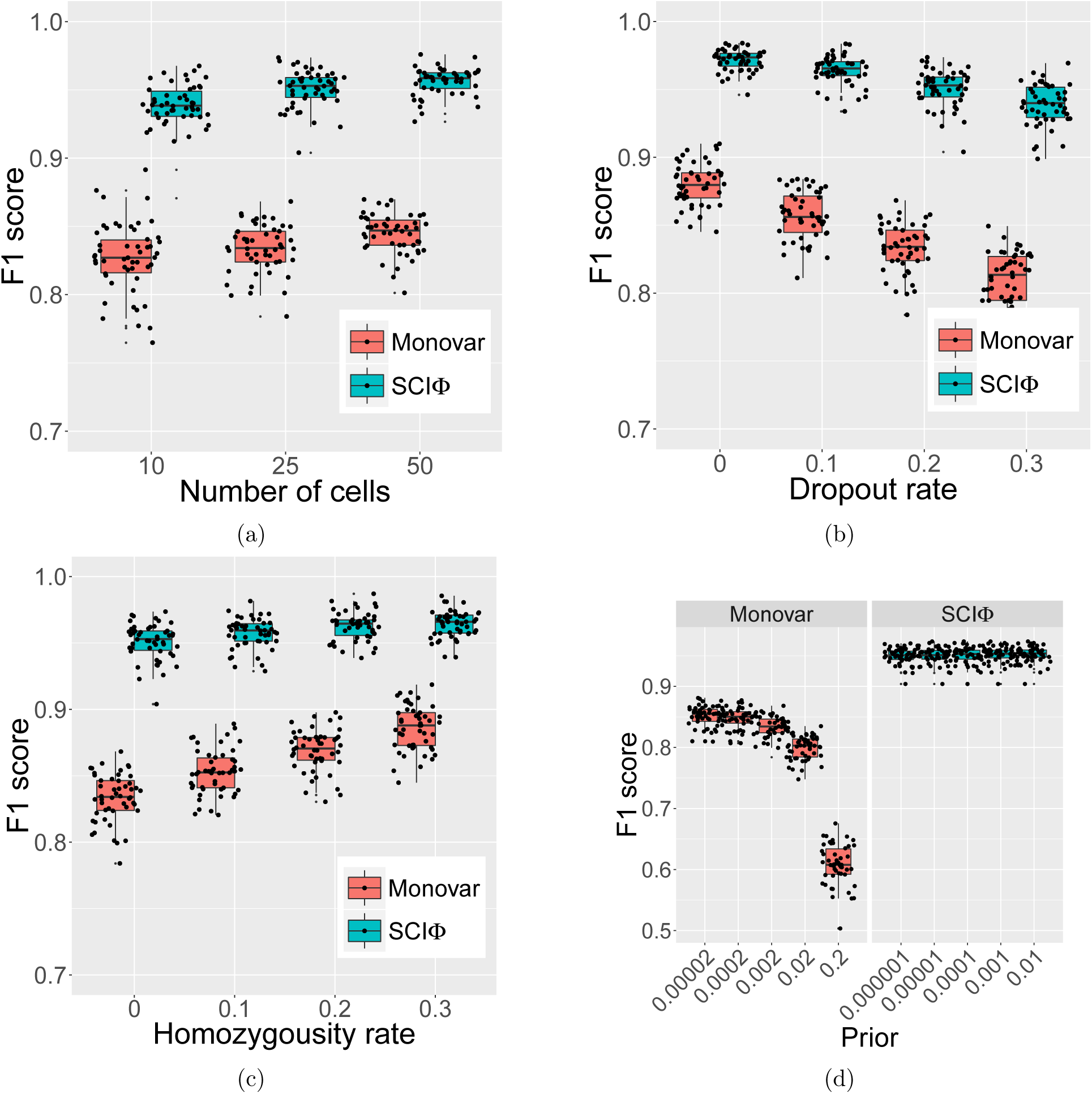
Summary statistics of the F1 performance from Monovar and SCIΦ on simulated data with *α* and *β* set to 1 for the Pólya urn MDA process. (a) F1 score depending on the number of cells. (b) F1 score depending on the drop-out rate. (c) F1 score depending on the homozygosity rate. (d) F1 score depending on the false positive prior for Monovar and the mutation prior for SCIΦ.

## References

[1] Christophe Andrieu and Johannes Thoms. A tutorial on adaptive MCMC. Statistics and Computing, 18:343–373, 2008.

[2] Rebecca A Burrell and Charles Swanton. Tumour heterogeneity and the evolution of polyclonal drug resistance. Molecular Oncology, 8:1095–1111, 2014.

[3] Frank B Dean, John R Nelson, Theresa L Giesler, and Roger S Lasken. Rapid amplification of plasmid and phage DNA using phi29 DNA polymerase and multiply-primed rolling circle amplification. Genome Research, 11:1095–1099, 2001.

[4] Xiao Dong, Lei Zhang, Brandon Milholland, Moonsook Lee, Alexander Y Maslov, Tao Wang, and Jan Vijg. Accurate identification of single-nucleotide variants in whole-genome-amplified single cells. Nature Methods, 14:491–493, 2017.

[5] Andreas D”oring, David Weese, T Rausch, and Knut Reinert. SeqAn an efficient, generic C++ library for sequence analysis. BMC Bioinformatics, 9(1):11, 2008.

[6] Charles Gawad, Winston Koh, and Stephen R Quake. Dissecting the clonal origins of childhood acute lymphoblastic leukemia by single-cell genomics. Proceedings of the National Academy of Sciences, 111:17947–17952, 2014.

[7] Moritz Gerstung, Christian Beisel, Markus Rechsteiner, Peter Wild, Peter Schraml, Holger Moch, and Niko Beerenwinkel. Reliable detection of subclonal single-nucleotide variants in tumour cell populations. Nature Communications, 3:811, 2012.

[8] Mel Greaves. Evolutionary determinants of cancer. Cancer Discovery, 5:806–820, 2015.

[9] Zuguang Gu, Roland Eils, and Matthias Schlesner. Complex heatmaps reveal patterns and correlations in multidimensional genomic data. Bioinformatics, 32(18):2847–2849, may 2016.

[10] Zheng Hu, Ruping Sun, and Christina Curtis. A population genetics perspective on the determinants of intra-tumor heterogeneity. BBA Reviews on Cancer, 1867:109–126, 2017.

[11] Katharina Jahn, Jack Kuipers, and Niko Beerenwinkel. Tree inference for single-cell data. Genome Biology, 17:86, 2016.

[12] Jack Kuipers, Katharina Jahn, and Niko Beerenwinkel. Advances in understanding tumour evolution through single-cell sequencing. BBA Reviews on Cancer, 1867:127–138, 2017.

[13] Jack Kuipers, Katharina Jahn, Benjamin J. Raphael, and Niko Beerenwinkel. Single-cell sequencing data reveal widespread recurrence and loss of mutational hits in the life histories of tumors. Genome Research, 27(11):1885–1894, oct 2017.

[14] Roger S Lasken. Genomic DNA amplification by the multiple displacement amplification (MDA) method. Biochemical Society Transactions, 37:450–453, 2009.

[15] Si Quang Le and Richard Durbin. SNP detection and genotyping from low-coverage sequencing data on multiple diploid samples. Genome Research, 21:952–960, 2011.

[16] Heng Li and Richard Durbin. Fast and accurate short read alignment with Burrows–Wheeler transform. Bioinformatics, 25:1754–1760, 2009.

[17] Heng Li, Bob Handsaker, Alec Wysoker, Tim Fennell, Jue Ruan, Nils Homer, Gabor Marth, Goncalo Abecasis, and Richard Durbin. The sequence alignment/map format and SAMtools. Bioinformatics, 25:2078–2079, 2009.

[18] Aaron McKenna, Matthew Hanna, Eric Banks, Andrey Sivachenko, Kristian Cibulskis, Andrew Kernytsky, Kiran Garimella, David Altshuler, Stacey Gabriel, Mark Daly, et al. The genome analysis toolkit: a MapReduce framework for analyzing next-generation DNA sequencing data. Genome Research, 20:1297–1303, 2010.

[19] Nicholas E Navin. Cancer genomics: one cell at a time. Genome Biology, 15:452, 2014.

[20] Nicholas E Navin. The first five years of single-cell cancer genomics and beyond. Genome Research, 25:1499–1507, 2015.

[21] R Development Core Team. R: A Language and Environment for Statistical Computing. R Foundation for Statistical Computing, Vienna, Austria, 2008. ISBN 3-900051-07-0.

[22] Jochen Singer, Hans-Joachim Ruscheweyh, Ariane L Hofmann, Thomas Thurnherr, Franziska Singer, Nora C Toussaint, Charlotte KY Ng, Salvatore Piscuoglio, Christian Beisel, Gerhard Christofori, et al. NGS-pipe: a flexible, easily extendable and highly configurable framework for NGS analysis. Bioinformatics, in press, 2017.

[23] Gregory R Smith and Marc R Birtwistle. A mechanistic beta-binomial probability model for mRNA sequencing data. PLoS ONE, 11:e0157828, 2016.

[24] Yong Wang, Jill Waters, Marco L Leung, Anna Unruh, Whijae Roh, Xiuqing Shi, Ken Chen, Paul Scheet, Selina Vattathil, Han Liang, et al. Clonal evolution in breast cancer revealed by single nucleus genome sequencing. Nature, 512:155–160, 2014.

[25] Hamim Zafar, Yong Wang, Luay Nakhleh, Nicholas Navin, and Ken Chen. Monovar: singlenucleotide variant detection in single cells. Nature Methods, 13:505–507, 2016.

